# Tumor-derived stearic acid induces macrophage Egr2 signaling to suppress anti-tumor immunity in breast cancer

**DOI:** 10.64898/2026.07.15.738781

**Authors:** Yongling Ning, Caijun Wu, Sailesh Phuyal, Xiaoling Hu, Hong Li, Robert A Mitchell, Jun Yan, Chuanlin Ding

**Author notes:** Correspondence: Dr. Chuanlin Ding, The Hiram C. Polk, Jr., MD Department of Surgery, UofL Health Brown Cancer Center, University of Louisville School of Medicine.; and Dr. Jun Yan, The Hiram C. Polk, Jr., MD Department of Surgery, UofL Health Brown Cancer Center, University of Louisville School of Medicine. The authors declare no potential conflicts of interest. Yongling Ning: The Second People’s Hospital of Changzhou, The Third Affiliated Hospital of Nanjing Medical University, Changzhou, 213003, China.

## Abstract

**Background:** Macrophages in breast cancer are highly heterogeneous and targeting both tissue-resident and bone marrow-derived macrophages represents a high-potential strategy in breast cancer immunotherapy.

**Methods:** Myeloid-specific Egr2 knockout (KO) mice were used to determine roles of macrophage Egr2 signaling in tumor progression and anti-tumor immune responses. RNA-seq analysis and Flow cytometry were used to examine regulation of Egr2 in tumor-associated macrophages (TAMs) and fatty acid-stimulated macrophages. Tumor cell and macrophage admixture study was used to evaluate effects of fatty acid educated macrophages on anti-tumor responses.

**Results:** Here we showed that early growth response 2 (Egr2) was highly expressed in mammary tissue macrophages and recruited macrophages upon tumor progression. Depletion of Egr2 in myeloid cells significantly decreased the immunosuppressive function of polarized M2-like macrophages and TAMs. Tumor progression was delayed in myeloid cell Egr2 knockout mice, which was associated with enhanced function of effector CD8^+^ T cells and NK cells. Mechanistically, we showed that retinol and nicotinamide metabolism in TAMs were impaired after Egr2 depletion. Further, tumor cell-derived stearic acid (SA) was identified as an important fatty acid inducing Egr2 expression. SA stimulation induced macrophage-mediated immunosuppression in an Egr2 dependent manner. Addition of oleic acid (OA) uniquely repressed SA-induced Egr2 expression and immunosuppressive activity in macrophages.

**Conclusions:** Our study thus uncovers a novel SA-Egr2 pathway driving macrophage immunosuppression and novel molecular mechanism of OA mediated anti-tumor effect.

**WHAT IS ALREADY KNOWN ON THIS TOPIC:** Lipid-associated macrophages (LAMs) accumulate lipids and promote immune suppression, tumor growth and metastasis in breast cancers. Fatty acids play a dual, complex role in tumor immunity, acting as both pro-tumorigenic and anti-tumorigenic effects. However, molecular mechanisms underlying fatty acid modulated macrophages and their roles in tumor remain not fully elucidated.

**WHAT THIS STUDY ADDS:** We demonstrate that Egr2 is essential for the immunosuppressive function of IL-4 polarized M2-like macrophages and TAMs. Tumor cell-derived stearic acid (SA) was identified as an important fatty acid inducing Egr2 expression and immunosuppression in macrophages. Adding oleic acid (OA) uniquely repressed SA-induced Egr2 expression and immunosuppression.

**HOW THIS STUDY MIGHT AFFECT RESEARCH, PRACTICE OR POLICY:** SA-Egr2 axis is a novel pathway driving macrophage immunosuppression. OA treatment enhances macrophage anti-tumor immunity via inhibiting Egr2 signaling and immunosuppression. The data supports the notion that diets enriched in OA may have potential for prevention and suppression of breast cancer.

## INTRODUCTION

Tumor-associated macrophages (TAMs) play essential roles at each stage of breast cancer progression and metastasis, and targeting TAMs is an attractive approach for cancer immunotherapy. However, the current strategies targeting macrophage-associated molecules show limited success. Targeting CCL2-CCR2 axis to inhibit macrophage recruitment is largely disappointing^1–4^. These can be explained in part by the fact that TAMs have dual origins and distinct functions. Previous studies have mainly focused on monocyte-derived macrophages (MDMs) due to the accumulation with an increased tumor burden. The roles of tissue-resident macrophages in breast cancer are not yet fully understood^4–7^. Mammary tissue macrophages (MTMs) are tissue-resident cells that display M2-like macrophage phenotype^4^. A recent study has demonstrated that MTMs contribute to early breast cancer development^8^. We postulate that targeting both tissue-resident and monocyte-derived macrophages provide a powerful approach to reprogram TAMs in breast cancer.

Lipid-associated macrophages (LAMs) have been identified in the tumor microenvironment of human breast cancer^9–11^. These LAMs can accumulate lipids and promote immune suppression, tumor growth and metastasis in breast cancers. Fatty acids (FAs) from tumor synthesis and diet play a dual, complex role in tumor immunity, acting as both pro-tumorigenic and anti-tumorigenic effects, which are dependent on the specific FA type and metabolism pathways in both cancer and immune cells^12–18^. However, molecular mechanisms underlying fatty acid modulated macrophages and their roles in tumor remain not fully elucidated.

Transcription factor early growth response 2 (Egr2) is a zinc finger transcription factor of the early growth response gene family. Previous data have demonstrated that Egr2 signaling in tumor-infiltrating T cells negatively regulates anti-tumor immunity^19–21^. Recent studies suggest that macrophage Egr2 signaling exhibits pro-tumor effect^22,23^. However, the function of Egr2 in macrophage heterogeneity in mammary gland and breast cancer are largely unexplored. Tumor factors inducing Egr2 expression and involvement of Egr2 in fatty acid modulated macrophages in tumor remain unknown. In this study, we demonstrate that Egr2 is essential for the immunosuppressive function of IL-4 polarized M2-like macrophages and TAMs. Further, tumor cell-derived stearic acid (SA) was identified as an important fatty acid inducing Egr2 expression and immunosuppression in macrophages. Adding oleic acid (OA) uniquely represses SA-induced Egr2 expression and immunosuppression. These findings support our hypothesis that SA-Egr2 axis is a novel pathway driving macrophage immunosuppression, and OA treatment enhances macrophage anti-tumor immunity via inhibiting Egr2 signaling and immunosuppression.

## MATERIALS AND METHODS

### Mice and tumor model

Myeloid-specific Egr2 knockout (KO) (LyzM^Cre^-Egr2^f/f^) and control mice (LyzM^wt^-Egr2^f/f^) were kindly provided by Dr. Thomas F. Gajewski. C57BL/6J mice, OVA TCR Tg OT-I and OT-II mice, transgenic FVB/MMTV-PyMT mice were purchased from the Jackson Laboratory. The murine mammary cancer cell lines E0771 and 4T1 cells were purchased from ATCC. The human breast cancer cell line MDA-MB-231 was kindly provided by Dr. Brian Clem. All cell lines were confirmed to be mycoplasma free and cultured in the completed DMEM medium containing 10% FBS. To establish tumor, 5×10^5^ E0771 tumor cells were mixed with Matrigel and injected into 4^th^ mammary fat pad of female C57BL/6J mice. Tumor sizes were measured twice a week with a caliper. The spontaneous breast tumors were harvested from 10-12 weeks old transgenic female FVB/MMTV-PyMT mice. Only female mice were used in tumor studies. All animals were maintained under specific pathogen-free conditions and handled in accordance with the protocols approved by The University Committee for Animal Welfare (UCAW) of University of Louisville.

### Preparation of single cell suspension, flow cytometry, and cell sorting

Mouse mammary glands and tumor tissues were digested with collagenase IV (300 U/ml) and DNase I (80 U/ml) in complete RPMI1640 medium for 30 min at 37°C. After incubation, the digestion was immediately stopped by addition of 5 ml cold medium. The cell suspension was then filtered through a 40 µm cell strainer and washed twice with complete medium. For flow cytometry analysis, the cells were blocked in the presence of anti-CD16/CD32 at 4°C for 10 min and stained on ice with the appropriate antibodies and isotype controls. For Egr2 expression measurement, cells were stained with surface markers followed by fixation and permeabilization using eBioscience Foxp3/Transcription Factor Staining Buffer Set. Cells were then stained with anti-mouse Egr2 antibody overnight at 4 °C. The samples were acquired using Cytek Aurora cytometry and analyzed using FlowJo software. The tumor-associated macrophages (TAMs, CD45^+^CD11b^high^F4/80^+^) were sorted by using BD FACSymphony™ S6 cell sorter. Fixable viability dye eFluor™ 780 (Thermo Fisher Scientific) was used in Flow cytometry analysis and cell sorting to exclude dead cells. A post sort analysis was performed to determine the purity of the macrophages with approximately 90% purity. Antibodies used in flow cytometry, cell sorting, and Western blot were listed in online supplementary table S1.

### Mass cytometry (CyTOF) and image mass cytometry (IMC) data acquisition and analysis

Tumor cells were stimulated with PMA plus Ionomycin in the presence of protein transport blocker Brefeldin A for 4 hours. The cell staining was performed according to the protocol of Maxpar Cell Surface Staining with Fresh Fix (Fluidigm). Data acquisition was performed on the CyTOF Helios system (Fluidigm). FCS files were normalized with Helios instrument work platform (FCS Processing) based on the calibration bead signal used to correct any variation in detector sensitivity. CyTOF data analysis was performed with FlowJo software. Total events were gated on CD45^+^ cells after removing beads, doublets, and dead cells. tSNE and FlowSOM clustering analysis for CyTOF data were performed using FlowJo Plugins platform. tSNE analysis was performed on all samples combined. Different immune populations were defined by the expression of specific surface and intracellular markers. For IMC on FFPE tissue sections, antigen retrieval was performed followed by staining with appropriate antibodies overnight at 4°C. Data acquisition was performed on Hyperion imaging mass cytometry. Antibodies used in CyTOF and IMC were listed in online supplementary table S2.

### Fatty acid analysis

Tumor cell conditioned media were collected from culture supernatants of E0771, 4T1, and MDA-MB-231 tumor cells (5 ×10^5^ in 4 ml culture medium for 3 days). TAMs were sorted from E0771 tumor-bearing control and Egr2 KO mice. The fatty acid composition and content in the conditioned media and TAMs were analyzed at the Lipidomics Core Facility of Wayne State University.

### In vitro BMDM culture, stimulation, and T cell immunosuppression assay

Bone Marrow-Derived Macrophages (BMDMs) are generated by culturing mouse bone marrow cells in the presence of M-CSF (20 ng/ml) for 6-7 days^24^. For M2 macrophage differentiation, BMDMs were stimulated with IL-4 and IL-13 (20 ng/ml) for 2 days. For fatty acid stimulation, BMDMs were cultured in the presence of BSA complex of 200 μM stearic acid (SA), palmitic acid (PA), oleic acid (OA), or BSA control for 48 hours. In some experiments, BMDMs were stimulated with 25 nM retinoic acid (RA) for 48 hours. The T cell immunosuppression assay was performed by coculture of M2 macrophages, fatty acid or RA-stimulated macrophages, or TAMs with CFSE labeled OT-II or OT-I mouse splenocytes in the presence of ovalbumin (OVA, 200 μg/ml for OT-II cells, 20 μg/ml for OT-I cells). T cell proliferation was evaluated by measuring CFSE dilution using Flow cytometry. The IFN-γ production by T cells after PMA/Ionomycin 4 to 6-hour stimulation was evaluated by intracellular cytokine staining and Flow cytometry.

### Quantitative real-time PCR

RNA extraction, cDNA synthesis, and quantitative real-time PCR reactions using SYBR Green Supermix (Bio-Rad) were performed as we reported ^25^. We normalized gene expression level to ribosomal protein L13a (RPL13A) housekeeping gene and represented data as fold differences by the 2^−ΔΔCt^ method. The primer sequences of real-time PCR were listed in online supplementary table S3.

### RNA sequencing and analysis

TAMs sorted from E0771 tumors and fatty acid 24-hour stimulated BMDMs were harvested for RNA extraction by using a QIAGEN RNAeasy Kit (QIAGEN). The quantity of purified RNA samples was measured by the RNA High Sensitivity Kit in the Qubit Fluorometric Quantification system (Thermo Fisher Scientific). Library preparation, sequencing, and analysis were performed by LC Sciences (https://lcsciences.com).

### Oxygen consumption rate (OCR) and extracellular acidification rate (ECAR) analysis

TAMs sorted from E0771 tumors were subjected to a Mito stress and glycolysis test to measure the bioenergetic function. Briefly, TAMs were washed twice with Agilent Seahorse XF Media supplemented with 1 mM pyruvate, 2 mM L-glutamine, and 10 mM D-glucose 1 h before the assay. A Seahorse XF Cell Mito Stress Test Kit (Agilent Technologies) was used with the final concentrations of 1.5 μM oligomycin, 2 μM carbonyl cyanide p-trifluoromethoxyphenylhydrazone (FCCP), and 0.5 μM rotenone/antimycin, respectively. For ECAR experiments, cells were washed twice with Agilent Seahorse XF Media supplemented with 2 mM L-glutamine 1 h before the assay, and then treated with 10 mM glucose, 1.0 μM oligomycin (Oligo), and 50 mM 2-Deoxy-D-glucose (2-DG).

### Tumor cell and macrophage admixture study

BMDMs were treated with 200 μM of SA, SA+OA, or BSA control for 48 hours. The admixture of E0771 tumor cells (2 × 10^5^/per mouse), BMDMs (2 × 10^5^/per mouse) and 50% Matrigel was injected into 4^th^ mammary fat pad of female C57BL/6J mice. Tumor progression was evaluated until day 17 post injection. For immune cell phenotyping, tumor tissues were collected on day 7 after injection.

### Analysis of clinical data sets

Egr2 gene expression in human macrophage subsets from breast cancer dataset GSE176078^26^ was performed by using Single Cell Portal (https://singlecell.broadinstitute.org/single_cell). Correlation between Egr2 and M2 macrophage infiltration or other genes were analyzed using TIMER3.0^27^, which incorporates 1086 BRCA samples. Triple negative breast cancer paraffin tissue sections were purchased from TissueArray.Com LLC (https://www.tissuearray.com).

### Statistical analysis

Data was analyzed using GraphPad Prism 10 software. An unpaired Student’s t-test and one-way or two-way ANOVA were used to calculate significance for two groups and three or more groups, respectively. All graph bars are expressed as mean ± SEM. Significance was assumed to be reached at p < 0.05. The p values were presented as follows: *p<0.05, **p<0.01, ***p<0.001, and ****p<0.0001.

## RESULTS

### Egr2 expression in macrophages

Previous studies have shown that Egr2 is a novel M2 macrophage marker and highly expressed in IL-4+IL-13 polarized M2 macrophages^28^. We further analyzed the data from The Immunological Genome Project (http://www.immgen.org) and found that Egr2 was highly expressed in tissue-resident MTMs, but not in bone marrow macrophages (BMMφ), splenic macrophages (SpMφ) and peritoneal macrophages (PecMφ) (figure 1B). Flow cytometry analysis confirmed that Egr2 was expressed in MTMs (CD45^+^F4/80^+^CD11b^+^) under steady-state conditions (figure 1C). Expression of Egr2 in M2 macrophages and MTMs was remarkably downregulated in conditional myeloid cell Egr2 KO (LyzM-cre-Egr2^f/f^) mice (figure 1A and C). Tissue-resident macrophages usually express low levels of CCR2 and MHC II compared to BM monocyte-derived macrophages. We examined Egr2 expressions in TAMs from MMTV-PyMT transgenic mice and found that Egr2 was expressed in both BM-derived CCR2^+^MHC II^high^ and tissue-resident CCR2^-^MHC II^low^ TAM subsets, but not neutrophils (figure 1D).

**Figure 1.**
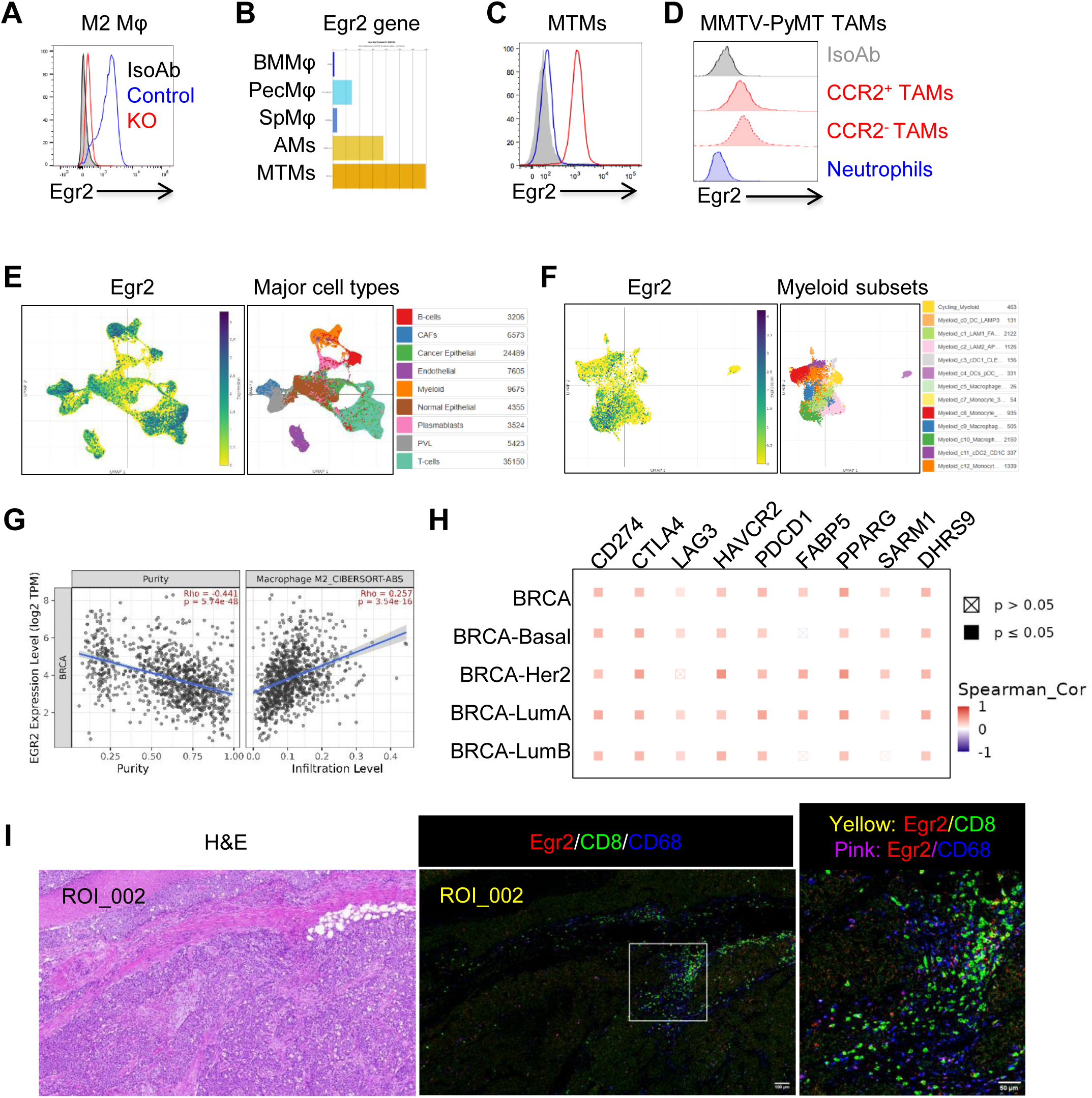
Egr2 expression in macrophages and correlation with immunosuppression. (A) Egr2 expression in control and Egr2 KO IL-4+IL-13 polarized M2 macrophages. (B) Egr2 mRNA levels in macrophages from different tissues. Data was from the Immunological Genome Project. (C) Egr2 expression in MTMs from control and Egr2 KO mice. (D) Egr2 expression in CCR2^+^ and CCR2^−^ TAMs and neutrophils from tumors of MMTV-PyMT mice. (E) ScRNA-seq analysis of Egr2 expression across major cell types. (F) ScRNA-seq analysis of Egr2 expression across myeloid cell subsets. Data of (E) and (F) was from breast cancer dataset GSE17607823 and re-analyzed by using Single Cell Portal. (G) Correlation of Egr2 expression with infiltration level of M2 macrophages in breast cancer patients. (H) Correlation of Egr2 expression and genes related to immunosuppression and cellular metabolism. Data of (G) and (H) was from the TIMER3.0 web platform (n = 1,086). (I) Expression of Egr2 in human macrophages from a TNBC patient. Tumor part in breast cancer tissue was verified by H&E staining (left). Macrophages, CD8 T cells, and Egr2 expression were revealed by using CD68 (blue), CD8 (green), and Egr2 (red) antibodies (right).

To demonstrate potential clinical significance, we analyzed Egr2 expression in human macrophages from single cell RNA sequencing dataset GSE176078, which identified six macrophage subsets from breast cancer patients including two clusters of lipid-associated macrophages (LAM1, C1 and LAM2, C2) and two “M2-like” clusters (C10, C5)^26^. Egr2 was highly expressed in myeloid cells, T cells, B cells, epithelial cells, and fibroblasts (figure 1E). In myeloid cells, Egr2 was highly expressed in M2-like macrophages (C10) and LAM1 (C1) clusters. By using publicly available web platform TIMER3.0 (https://compbio.cn/timer3), we further analyzed the significance of macrophage Egr2 expression in breast cancer patients (n=1,086). As shown in figure 1G, Egr2 expression was correlated with M2 macrophage infiltration levels. Egr2 expression was also positively correlated with genes related to immunosuppression (*CD274, CTLA4, LAG3, HAVCR2, PDCD1*) and cellular metabolism (*FABP5, PPARG, SARM1, DHRS9*) (figure 1H). We further used Imaging Mass Cytometry to verify macrophage Egr2 expression and interaction between Egr2^+^ macrophages and CD8^+^ T cells. As shown in figure 1I, Egr2 was expressed in both macrophages and CD8^+^ T cells. Importantly, Egr2^+^ macrophages conjugated to CD8^+^ T cells to form cluster within tumor tissue of triple negative breast cancer patients, indicating that these macrophages may directly dictate anti-tumor outcomes of CD8^+^ T cells. Together, these data suggest that Egr2 is a novel and important molecule which may contribute to macrophage immunosuppression and metabolism in the tumor microenvironment of breast cancer.

### Egr2 regulates immunosuppression of M2 macrophages and tumor educated macrophages

Egr2 expression is significantly upregulated during M2 macrophage polarization^28,29^. Whether Egr2 regulates M2 macrophage immunosuppression remains unknown. We next sought to determine the functional importance of Egr2 in IL-4+IL-13 polarized M2 macrophages. As shown in figure 2A and 2B, M2 macrophages exerted potent immunosuppressive activity for transgenic OVA-specific OT-II (CD4^+^) and OT-I (CD8^+^) T cell activation. Notably, the immunosuppressive activity of M2 macrophages for OVA-induced CD4 and CD8 T cell proliferation was significantly reduced in Egr2 KO macrophages (figure 2A and 2B). The expression of M2 associated genes (*CEBPB, PPARG, and CHIL3*) were significantly decreased in Egr2 KO M2 macrophages. Other M2 genes, such as *ARG1 and RETNLA (FIZZ1),* were not altered after Egr2 depletion (figure 2C). Western blotting further confirmed that PPARγ was remarkably reduced in Egr2 KO M2 macrophages (figure 2D). To reveal whether down-regulation of PPARγ is responsible for Egr2 deficiency-mediated immunosuppression reversing in M2 macrophages, GW9662, a selective irreversible PPARγ antagonist, was used in M2 macrophage differentiation. As shown in figure 2E, PPARγ inhibition significantly reversed immunosuppression in control M2 macrophages, but not in Egr2-deficient M2 macrophages.

**Figure 2.**
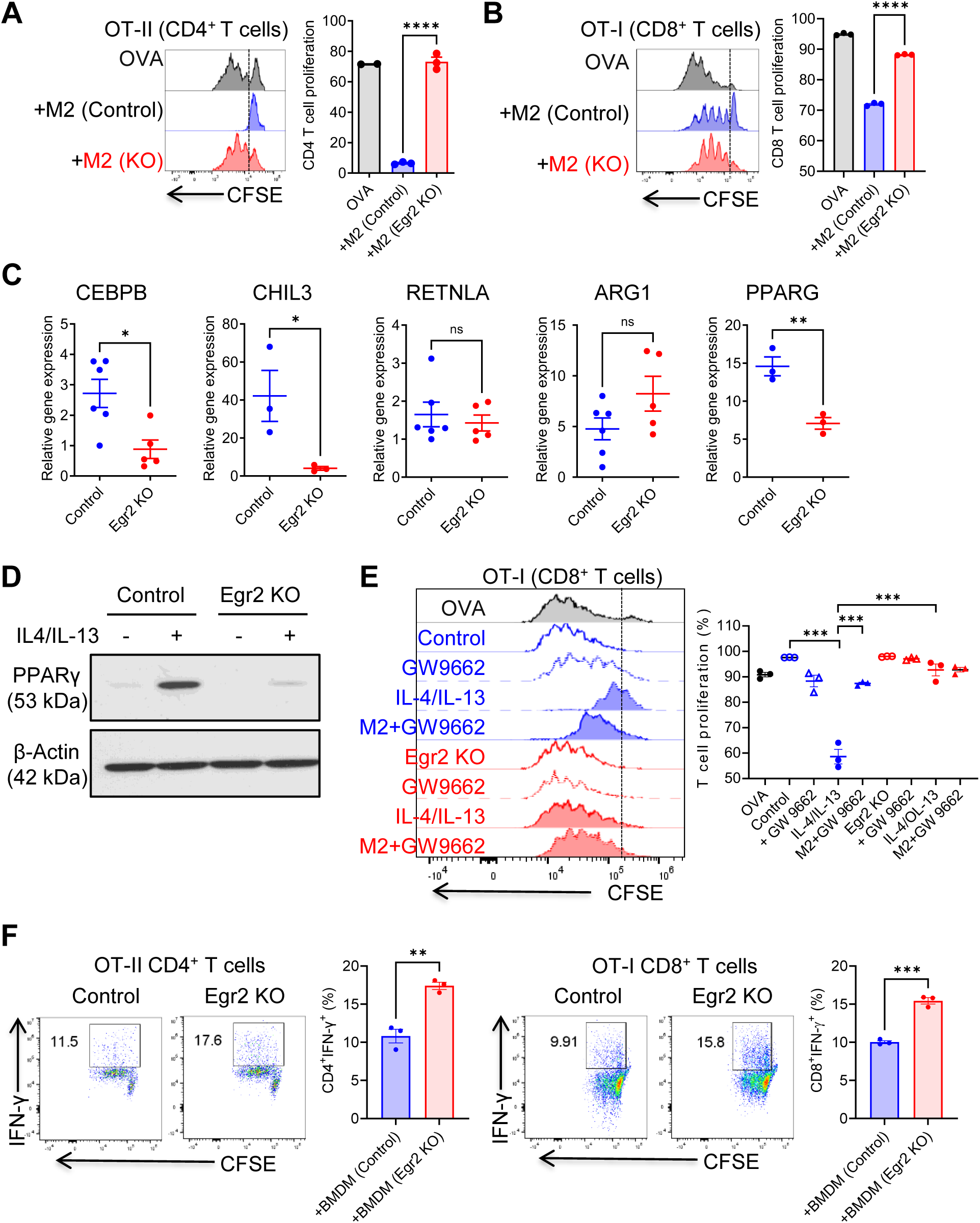
Egr2 regulates macrophage immunosuppression. (A and B) Control and Egr2 KO IL-4+IL-13 polarized M2 macrophages were co-cultured with CFSE labeled OT-II or OT-I splenocytes in the presence of OVA (200 μg/ml and 20 μg/ml, respectively) for 3 days. Proliferation of CD4^+^ T cells (A) and CD8^+^ T cells (B) was determined by measuring CFSE dilution. (C) Gene expression in the control and Egr2 KO IL-4+IL-13 polarized M2 macrophages were determined by using qRT-PCR. Each dot represents an individual mouse. (D) PPARγ and β-actin expression in the control and Egr2 KO IL-4+IL-13 polarized M2 macrophages were determined by using Western blot. (E) Control and Egr2 KO BMDMs were cultured in the presence of IL-4+IL-13 with or without GW9662 (10 μM) for 2 days and then co-cultured with OT-I splenocytes. CD8^+^ T cell proliferation was determined by measuring CFSE dilution. Summarized data was shown. (F) Control and Egr2 KO BMDMs were cultured in the presence of 25% E0771 conditioned medium for 2 days and then co-cultured with OT-II and OT-I splenocytes for 2 days. IFN-γ production by CD4^+^ and CD8^+^ T cells after PMA/Ionomycin 4 to 6-hour stimulation was evaluated by intracellular cytokine staining and Flow cytometry. Summarized data was shown. *p<0.05; **p<0.01; ***p<0.001; ****p<0.0001.

To demonstrate the potential of macrophage Egr2 signaling in tumor-induced immunosuppression, bone-marrow-derived macrophages (BMDMs) were cultured in the presence of tumor cell conditioned media for two days. T cell immunosuppression assay showed that IFN-γ production by CD8 and CD4 T cells was significantly increased in the coculture with Egr2-deficient macrophages compared to control macrophages (figure 2F). Together, these data demonstrate that Egr2 is a critical regulator for the immunosuppressive function of M2 macrophages and tumor educated macrophages.

### Macrophage intrinsic Egr2 negatively regulates anti-tumor immunity

To determine the significance of macrophage Egr2 signaling in tumor, we established orthotopic E0771 breast cancer model by injection of E0771 cells into 4^th^ mammary fat pad of female mice. A significant delay in tumor growth was observed in the conditional myeloid cell Egr2 KO mice compared to that in the control mice (figure 3A and B). Next, we used mass cytometry (CyTOF) to determine the effect of myeloid cell Egr2 deficiency on immune cell landscape within tumor microenvironment after *ex vivo* PMA/Ionomycin 4 to 6-hour stimulation in the presence of brefeldin A. FlowSOM analysis identified a total of 20 immune cell subtypes, including myeloid cells (population 8, 9, 11, and 13), CD4 T cells (population 2 and 3), CD8 T cells (population 4, 5, and 6), NK cells (population 7), B cells (population 0), and DCs (population 10) (figure 3C). Among them, Ly6C^+^CD8^+^ T cells and NK cells were increased in Egr2 KO mice compared to that in the control mice (figure 3D). Flow cytometry analysis showed that CD8^+^ T cells (figure 3E), especially IFN-γ^+^TNF-α^+^ effector CD8^+^ T cells (figure 3F), were significantly increased in the Egr2 KO mice. It has been reported that Ly6C^+^CD8^+^ T cells are more effective in tumor growth suppression^12,30^. We further examined Ly6C^+^CD8^+^ T cell subset and found that Egr2 deficiency resulted in an expansion of Ly6C^+^CD8^+^ T cells (figure 3G). Compared to Ly6C^−^CD8^+^ T cells, Ly6C^+^CD8^+^ T cells produced higher levels of IFN-γ^+^ and TNF-α^+^ (figure 3H). Additionally, NK cell expansion in Egr2 KO mice was revealed by Flow cytometry (figure 3I). These NK cells produced higher level of IFN-γ (figure 3J). Together, these data demonstrate that targeting myeloid cell Egr2 signaling improves anti-tumor immunity in breast cancer.

**Figure 3.**
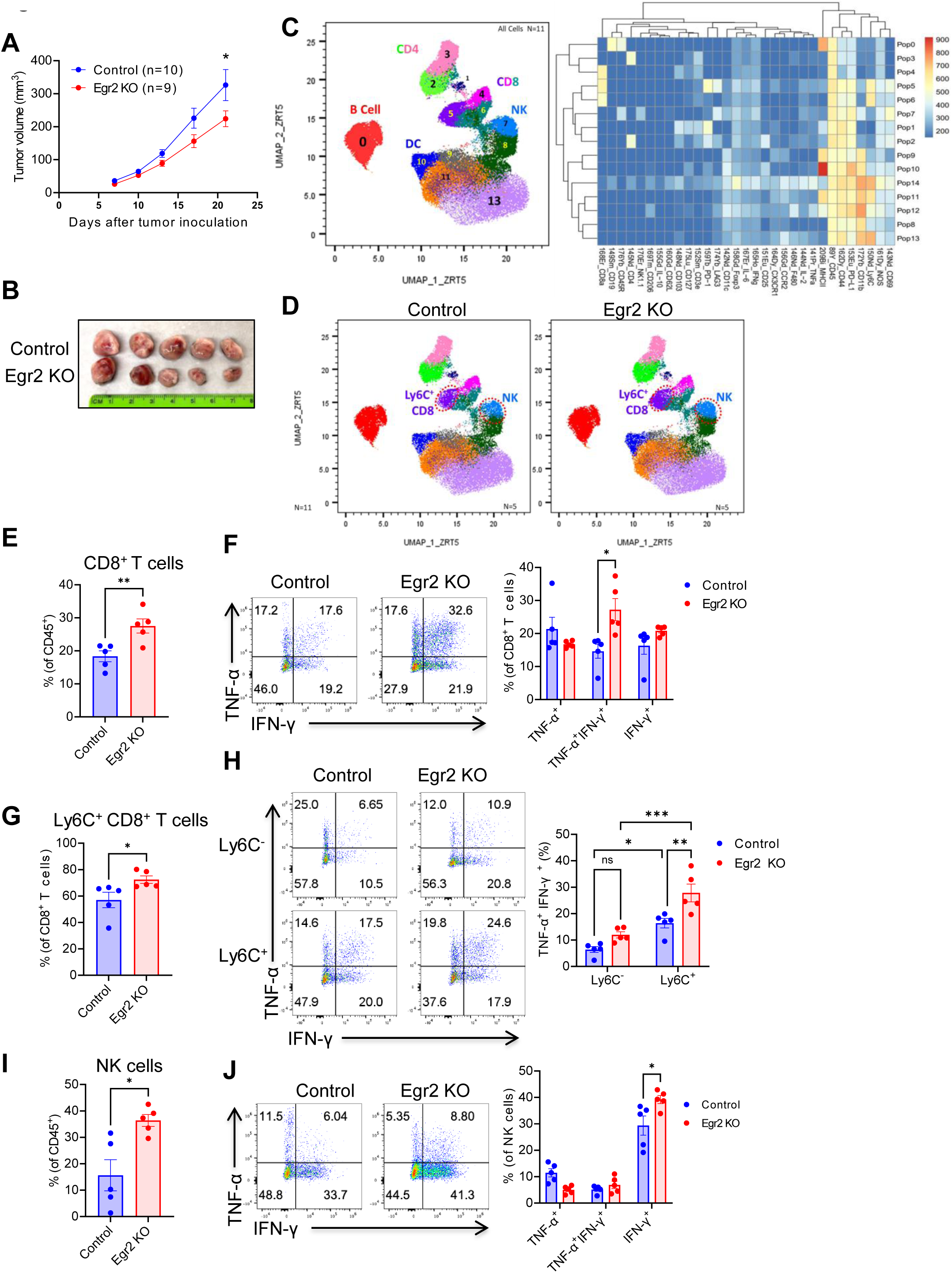
Macrophage intrinsic Egr2 negatively regulates anti-tumor immunity. (A and B) 5×10^5^ E0771 tumor cells were mixed with Matrigel and injected into 4^th^ mammary fat pad of female control and Egr2 KO mice. Tumor sizes were measured twice a week with a caliper. The tumor volumes (A) were summarized using calculation formula (wide^2^ × length)/2. The representative tumors from control and Egr2 KO mice at day 21 were shown (B). (C) viSNE analysis of CyTOF immunophenotyping of tumors from E0771 tumor-bearing control and c-Maf KO mice, all samples combined (n=11, left). FlowSOM clustering into 20 final immune cell types showing as a normalized expression heatmap (right). (D) viSNE analysis of CyTOF showing different Ly6C^+^CD8^+^ T cells and NK cell clusters in tumors from control and c-Maf KO mice (n=5). (E) Percentages of CD8^+^ T cells within CD45^+^ leukocytes. (F) IFN-γ and TNF-α producing CD8^+^ T cells. (G) Percentages of Ly6C^+^ cells within CD8^+^ T cells. (H) Comparison of IFN-γ and TNF-α production in Ly6C^−^ and Ly6C^+^ CD8^+^ T cells from control and Egr2 KO mice. (I) Percentages of NK cells within CD45^+^ leukocytes. (J) IFN-γ and TNF-α producing NK cells. Each dot represents an individual mouse. *p<0.05; **p<0.01; ***p<0.001.

### Egr2 regulates retinol and nicotinamide metabolism in tumor-associated macrophages

To understand how macrophage Egr2 signaling regulates anti-tumor immune responses, TAMs from E0771 tumor-bearing control and Egr2 KO mice were co-cultured with OVA TCR transgenic OT-I and OT-II splenocytes. As shown in figure 4A, both CD8^+^ and CD4^+^ T cells produced higher IFN-γ when co-culturing with Egr2 deficient TAMs compared to control TAMs. Next, we performed RNA-seq analysis to determine how Egr2 signaling regulates macrophage immunosuppressive activity. Altogether, 30 genes showed up- and downregulation. Among these genes, *Sarm1* and *Dhrs9* were significantly downregulated in Egr2 deficient TAMs compared to control TAMs (figure 4B). Gene set enrichment analysis showed that both retinol metabolism and nicotinate-nicotinamide metabolism were negatively enriched in Egr2 deficient TAMs (figure 4C). Dhrs9 is a stable marker for human regulatory macrophages^31^ and contributes to regulation of retinol metabolism. A prior study showed that retinoic acid regulated monocyte differentiation and promoted immune suppression in tumor^32^. To determine whether Egr2 plays an important role in regulation of retinol metabolism, BMDMs from control and Egr2 KO mice were cultured in the presence of retinoic acid (RA) and co-cultured with OT-I splenocytes. As shown in figure 4D, retinoic acid treated macrophages inhibited IFN-γ production in CD8^+^ T cells, which was abrogated in the Egr2 deficient macrophages. Sarm1 is a critical metabolic sensor and executioner enzyme in nicotinate and nicotinamide metabolism^33^. Downregulation of the nicotinate and nicotinamide metabolism (also known as the NAD^+^ salvage pathway) directly reduces cellular NAD^+^ levels inducing a metabolic shift towards glycolysis^34^. To determine how Egr2 regulates and maintains cellular energy metabolism, TAMs from control and Egr2 KO mice were sorted for ECAR and OCR Seahorse assay. We observed an increase in glycolysis in Egr2 deficient TAMs compared to control TAMs (figure 4E) whereas no significant changes in oxidative phosphorylation measured as OCR (online supplemental figure S1). Together, these data suggest that Egr2 plays important role in regulating TAM metabolism linked to their phenotype in anti-tumor responses.

**Figure 4.**
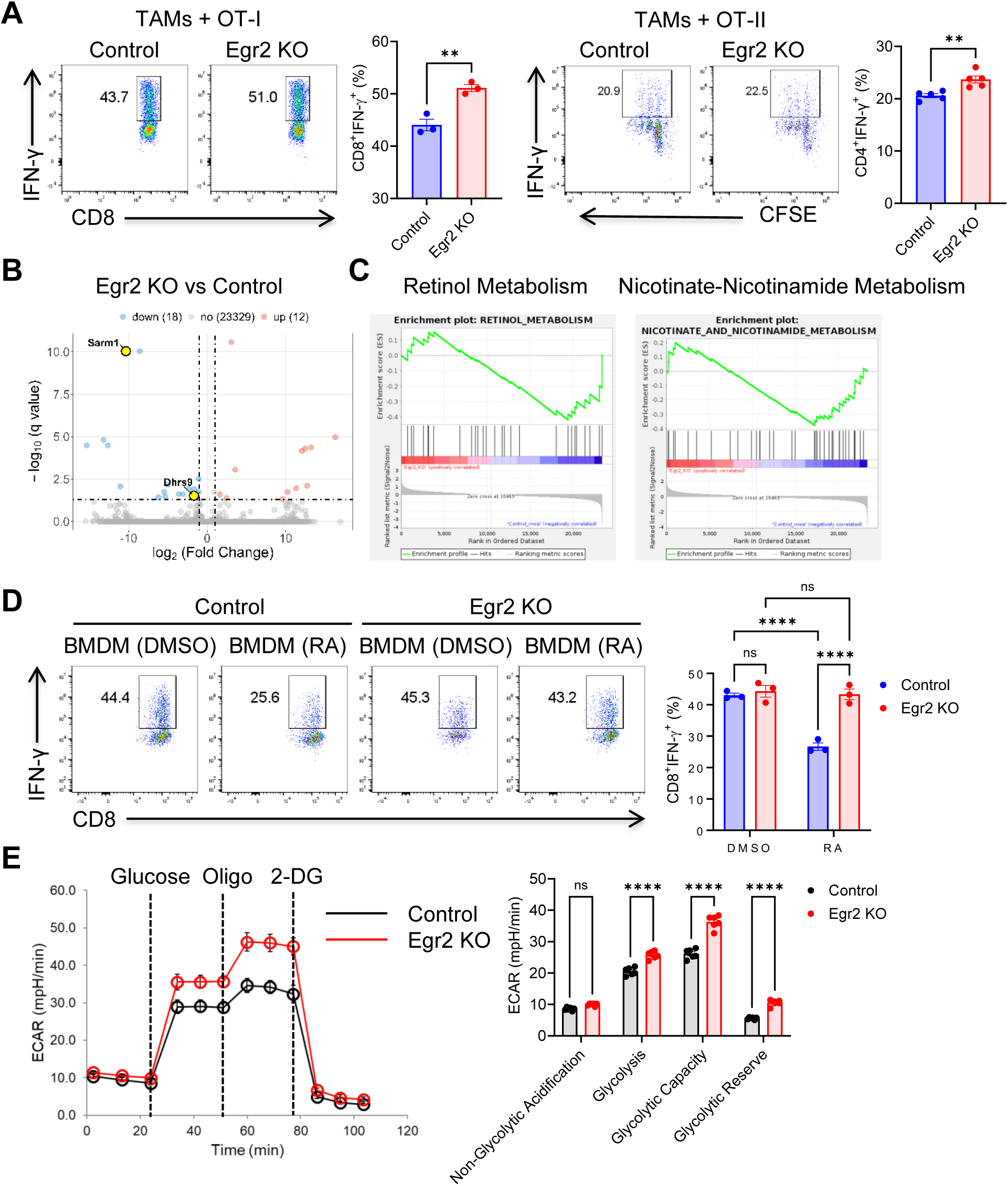
Egr2 regulates retinol and nicotinamide metabolism in TAMs. (A) Sorted TAMs from E0771 tumors were co-cultured with OT-I and OT-II splenocytes for 2 days. IFN-γ production by CD4^+^ and CD8^+^ T cells after PMA/Ionomycin 4 to 6-hour stimulation was evaluated by intracellular cytokine staining and Flow cytometry. Summarized data was shown. (B) Volcano plot showing fold change (FC) and *p* value for the comparison of TAMs from E0771 tumors of control and Egr2 KO mice (n = 3). (C) Gene set enrichment analysis showing retinol metabolism and nicotinate-nicotinamide metabolism in Egr2 deficient TAMs compared to control TAMs. (D) BMDMs were cultured in the presence of RA (25 nM) for 2 days and co-cultured with OT-I splenocytes for 2 days. IFN-γ production by CD8^+^ T cells after PMA/Ionomycin 4 to 6-hour stimulation was evaluated and summarized. (E) Seahorse glycolysis test in TAMs from E0771 tumors of control and Egr2 KO mice. Summary of relative values of ECAR bioenergenic profiling was shown. **p<0.01; ****p<0.0001.

### Tumor-derived stearic acid (SA) promotes immunosuppression of macrophages via regulation of Egr2

Fatty acid metabolism in immune cells in the tumor microenvironment is a new target of tumor immunotherapy^13^. A recent study showed that tumor-derived arachidonic acid (AA) reprogramd neutrophils to promote immune suppression in triple-negative breast cancer^17^. Gene set enrichment analysis of TAMs from E0771 tumors revealed downregulated signatures related to fatty acid synthesis and degradation pathway in the Egr2 KO TAMs (figure 5A). Several fatty acid degradation related genes, such as *Hadh, Hadhb, Adh5, Acat1, Acaa2, Acadm, and Aldh3a2 were* downregulated in Egr2 KO TAMs (online supplemental figure S2A). To determine whether de novo fatty acid synthesis contributes to pro-tumor effect of Egr2^+^ macrophages, TAMs were sorted from E0771 tumors for fatty acid profiling. As shown in online supplemental figure 2B, higher levels of linoleic acid and arachidonic acid were observed in TAMs, but there was no difference between control and Egr2-deficient TAMs.

**Figure 5.**
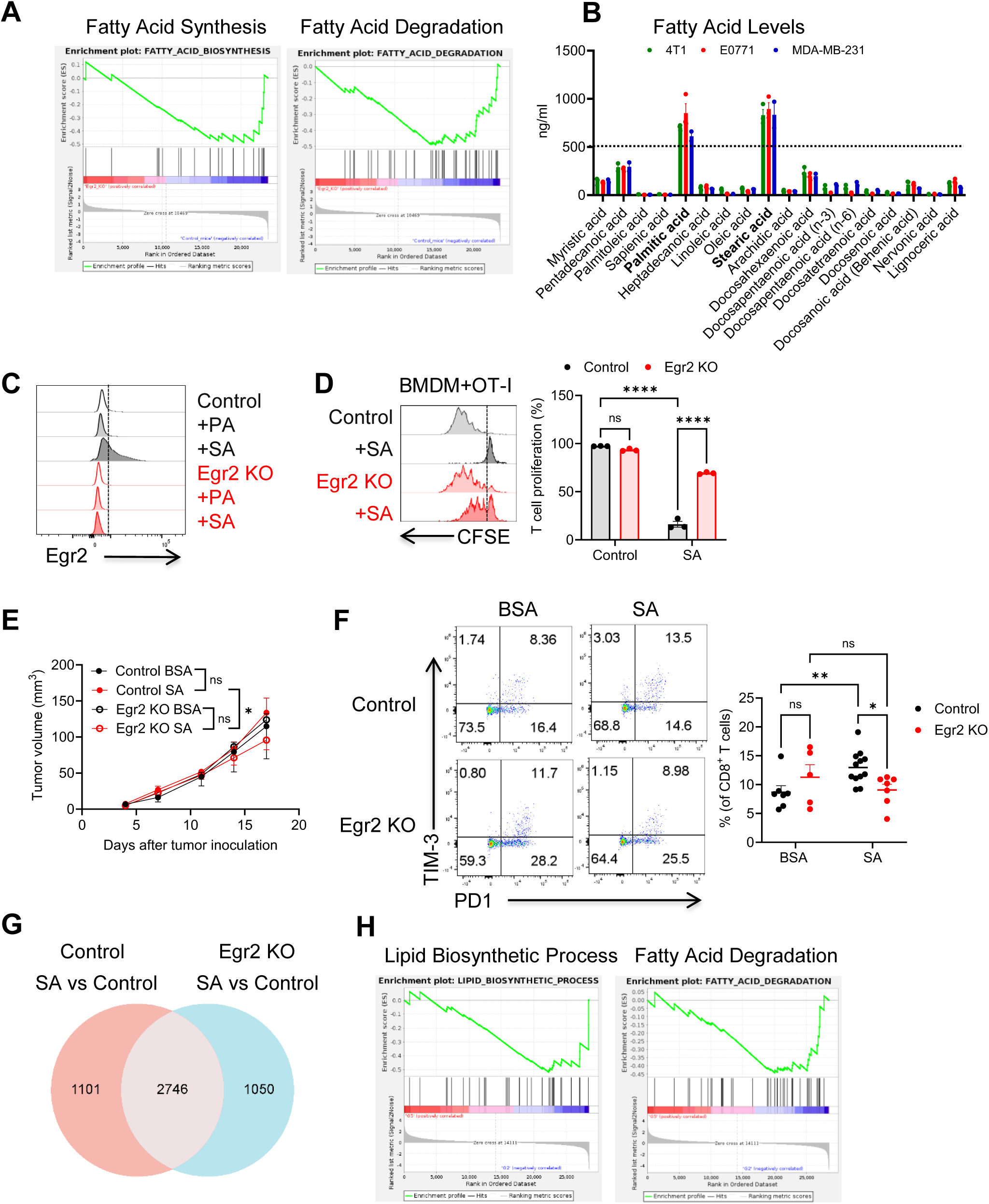
Stearic acid (SA) promotes immunosuppression of macrophages via regulation of Egr2. (A) Gene set enrichment analysis showing fatty acid synthesis and fatty acid degradation in Egr2 deficient TAMs compared to control TAMs. (B) Contents and levels of fatty acid in the conditioned media of 4T1, E0771, and MDA-MB-231 cells. (C) Egr2 expression in BMDMs from control and Egr2 KO mice after 48-hours stimulation with 200 μM PA, SA, or BSA control. (D) BMDMs from control and Egr2 KO mice were cultured with 200 μM SA for 48 hours and then co-cultured with OT-I splenocytes for 3 days. T cell proliferation was determined by measuring CFSE dilution. (E) Tumor development after injection of mixture of E0771 cells and SA or BSA treated control and Egr2 KO BMDMs. (F) Percentages of PD1^+^TIM-3^+^ exhausted CD8^+^ T cells on day 7 in tumor cell and macrophage admix experiments. Each dot represents an individual mouse. (G) Venn diagram showing common and unique genes after SA stimulation in control and Egr2 KO BMDMs (n=3) for 24 hours. (H) Gene set enrichment analysis showing lipid biosynthetic process and fatty acid degradation in SA treated Egr2 deficient BMDMs compared to control BMDMs. *p<0.05; **p<0.01; ****p<0.0001.

To identify tumor cell-derived factors driving Egr2 expression, conditioned media from breast cancer cell lines (mouse 4T1, E0771, and human MDA-MB-231) were collected for fatty acid analysis. Higher levels of stearic acid (SA) and palmitic acid (PA) were identified whereas OA levels were significantly low (figure 5B). Flow cytometry analysis revealed that treatment of SA, but not PA, induced Egr2 expression in macrophages (figure 5C). To investigate the effect of SA on macrophages, BMDMs from control and Egr2 KO mice were cultured in the presence of BSA-SA complex or BSA control for 2 days followed by coculture with CFSE labeled OT-I splenocytes. As shown in figure 5D, SA treated macrophages exhibited potent immunosuppression for T cell activation. This immunosuppression was substantially reduced in the Egr2 KO macrophages.

To investigate the contribution of SA educated macrophages to tumor progression, tumor cell and macrophage admix experiments were performed by injection of mixture of E0771 tumor cells and SA educated macrophages. Compared to BSA educated macrophages, SA educated macrophages promoted a trend increase of E0771 tumor progression. The pro-tumor effect of SA educated macrophages was significantly reduced in the SA educated Egr2 KO macrophages (figure 5E). Although there was no difference of IFN-γ^+^TNF-α^+^ producing CD8^+^ T cells among groups of mice (online supplemental figure S3A), PD-1^+^TIM-3^+^ exhausted CD8^+^ T cells were increased in the tumors from SA educated control macrophages and significantly decreased in the tumors from SA educated Egr2 KO macrophages (figure 5F).

To understand how Egr2 contributes into SA-induced macrophage phenotypic changes, BMDMs from control and Egr2 KO mice were stimulated with SA for 24 hours followed by RNA-seq analysis. A comparison of differentially expressed genes among different groups was illustrated in a Venn diagram (figure 5G). Among these genes, 1101 genes were identified as differentially expressed in SA-treated control macrophages but not in SA-treated Egr2 KO macrophages. Interestingly, downregulation of lipid biosynthetic process and fatty acid degradation pathways were revealed in SA-treated Egr2 KO macrophages compared to the SA- treated control macrophages (figure 5H). These data suggest that SA-Egr2 pathway promotes macrophage immunosuppression via regulation of lipid metabolism.

### Oleic acid (OA) reverses SA-induced Egr2 expression and immunosuppression

OA is the most abundant monounsaturated fatty acid in the human diet and highly enriched in olive oil. Studies have suggested that OA has potential anti-tumor activity^35,36^. A recent study showed that OA restored the impaired antitumor immunity of γδ-T cells induced by PA^37^. However, a prior study showed that pretreatment of thioglycolate-elicited peritoneal macrophages with OA before inducing M2 polarization increased some M2 macrophage markers^38^. To understand whether OA treatment regulates macrophage function, BMDMs were treated with SA, SA+OA, and control BSA for 24 hours. RNA-seq analysis revealed that SA-induced Egr2 expression was reversed after addition of OA (figure 6A). This data was verified by Flow cytometry analysis (figure 6C) and consistent with the finding that Egr2 drives the differentiation of Ly6C^high^ monocytes into fibrosis-promoting macrophages^39^. Gene sets related to antigen-presenting (*Ciita, H2-DMb2, H2-DMa, H2-DMb1, and H2-Ab1*), immunosuppression (*Ikbkg, CD274, Nfkbib, Lat, Ticam1*), and retinol metabolism (*Rdh13, Dhrs9, Ugt1a6a, Adh7*) were differently expressed in the SA or OA+SA treated macrophages (figure 6A). Additionally, antigen-presenting pathway was significantly downregulated in SA-treated macrophages and upregulated after addition of OA (figure 6B), which was consistent with MHC class II expression in SA or SA+OA stimulated macrophages examined by Flow cytometry (figure 6C). In contrast, retinol metabolism gene pathway was significantly upregulated in SA-treated macrophages and downregulated after addition of OA (figure 6B).

**Figure 6.**
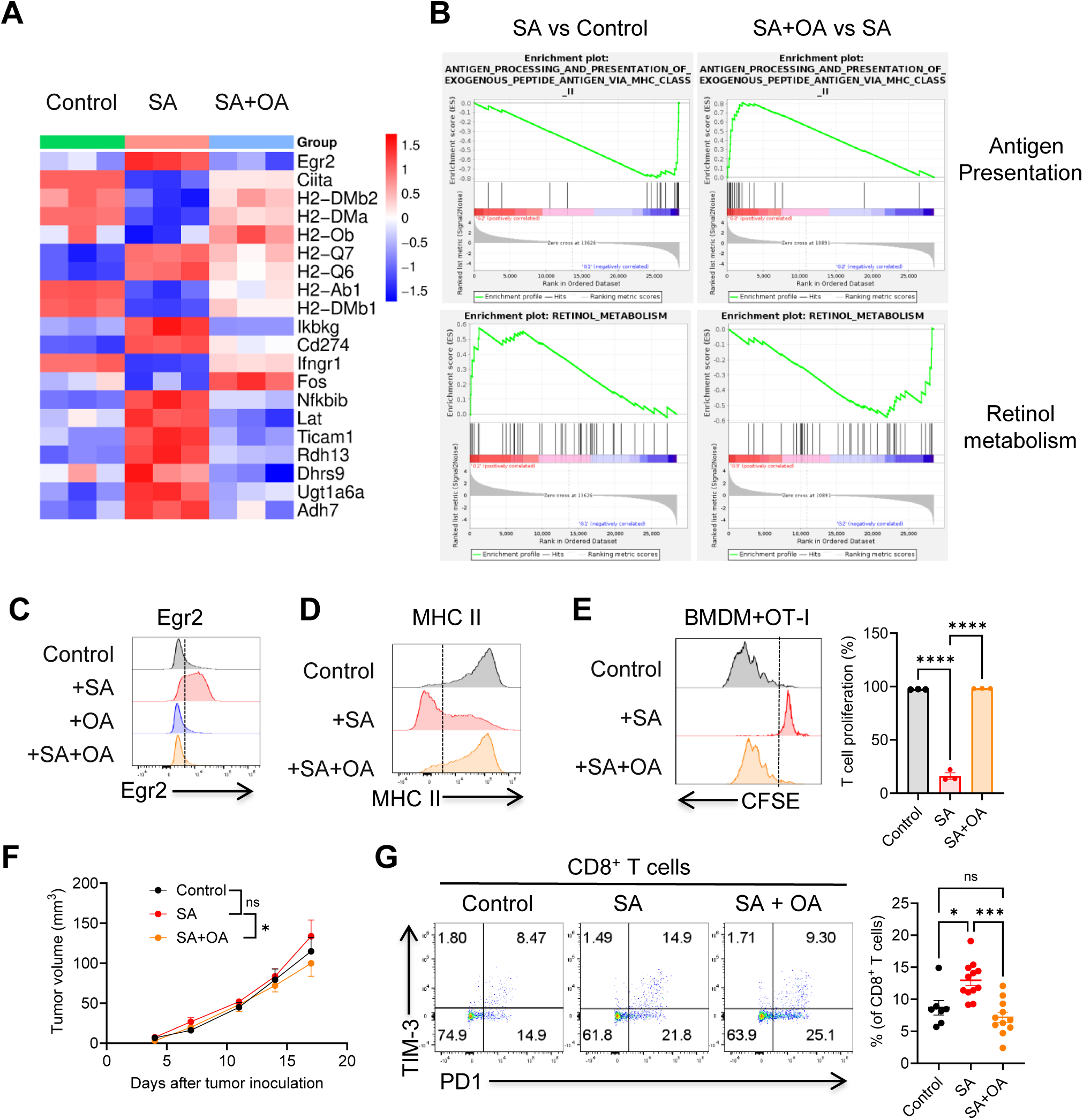
OA reverses SA-induced Egr2 expression and immunosuppression. (A) Heatmap showing the clustering of antigen presentation, immunosuppression, and retinol metabolism-related genes in three groups (n=3) based on log-relative abundances. (B) Gene set enrichment analysis showing antigen presentation and retinol metabolism in SA or SA+OA treated BMDMs. (C) Egr2 expression in BMDMs after stimulation with SA or SA+OA. (D) MHC class II expression in BMDMs after stimulation with SA or SA+OA. (E). BMDMs were cultured with 200 μM SA or SA+OA for 48 hours and then co-cultured with OT-I splenocytes for 3 days. T cell proliferation was determined by measuring CFSE dilution. (F) Tumor development after injection of mixture of E0771 cells and SA or SA+OA treated BMDMs. (F) Percentages of PD1^+^TIM-3^+^ exhausted CD8^+^ T cells on day 7 in tumor cell and macrophage admix experiments. Each dot represents an individual mouse. *p<0.05; ***p<0.001; ****p<0.0001.

To examine macrophage functional changes after OA treatment, BMDMs were treated with SA and SA+OA for 2 days followed by coculture with CFSE labeled OT-I splenocytes. T cell proliferation was significantly inhibited after coculture with SA-treated macrophages, which was remarkably reversed in the co-culture with SA+OA treated macrophages (figure 6E). We further performed tumor cell and macrophage admix study to evaluate the effects of OA educated macrophage on tumor progression. As shown in figure 6F, addition of OA reversed pro-tumor effect of SA educated macrophages. Increase of PD-1^+^TIM-3^+^ exhausted CD8 T cells in tumors injected with SA-treated macrophages was reversed in the tumors injected with SA+OA treated macrophages (figure 6G), although there was no change of IFN-γ^+^TNF-α^+^ effector CD8 T cells among groups of mice (online supplemental figure S3B). This data suggests that not only the sources but also the balance of fatty acids play critical roles in regulating macrophage function in tumor. Incorporating OA enriched diets may benefit breast cancer patients by reversing Egr2-mediated macrophage immunosuppression.

## DISCUSSION

Identifying novel immunological targets in tissue-macrophages for controlling early-stage breast cancer formation is essential. Targeting both tissue-resident macrophages and recruited monocyte-derived macrophages can elicit powerful anti-tumor immunity in breast cancer. Egr2 signaling in tumor-infiltrating T cells negatively regulates anti-tumor immunity^19–21^. The current study further reveals that Egr2 is expressed in both mammary tissue resident and BM monocyte-derived macrophages. Egr2 expression is associated with immunosuppression of macrophages. Targeting myeloid cell Egr2 can significantly enhance anti-tumor immunity in breast cancer.

The tumor microenvironment is rich in a diverse array of fatty acids and other lipids. Lipid-associated macrophages have been isolated in the tumor microenvironment of mouse tumor models and human breast cancer^9,10,26^. Targeting macrophage lipid metabolism is a promising strategy for tumor immunotherapy^9,10,12,40^. Studies also demonstrate that the type and composition of dietary fats can influence tumor growth and the effectiveness of anti-tumor immune responses^41^. A recent study discovered that tumor-derived arachidonic acid (AA) reprograms neutrophils to promote immune suppression^17^. Although previous studies have shown that Egr2 regulates lipid biosynthesis in peripheral nerve myelination^42^ and steatohepatitis^43^, the roles of Egr2 in regulating macrophage fatty acid metabolism in tumor remains unknown. PPARγ is crucial for fatty acid metabolic reprogramming and fatty acid oxidation (FAO) in T cells^44,45^. The current study demonstrates that IL-4 induced PPARγ expression is controlled by Egr2 and responsible for Egr2 mediated M2-like macrophage immunosuppressive activity. We further demonstrate a previously unidentified mechanism by which tumor-derived SA induces Egr2 expression and immunosuppression in macrophages. Egr2 signaling can regulate fatty acid metabolism in TAMs.

Obesity is associated with an increased risk of developing and worse prognosis in breast cancer patients. It is known that obesity reprograms mammary adipose tissue macrophages that support breast cancer formation^46^. In addition to the tumor microenvironment, SA is also found in various foods, such as beef fat, pork fat, cocoa butter, and processed foods. A recent study showed that the source of dietary fat influences anti-tumor immunity in obese mice^41^. Specifically, high-fat diets derived from lard, beef tallow or butter, but not from coconut oil, palm oil or olive oil, accelerate tumor growth. SA is significantly higher in butter high-fat diet^41^. SA associated metabolite stearoyl-carnitine (CAR18:0) is increased in the butter enriched high-fat-diet and directly inhibits CD8 T cell cytotoxicity^41^. These data strongly suggest that SA is an important pro-tumor fatty acid in the tumor microenvironment and high-fat diet induced obese. The current study further demonstrates that SA-induced macrophage immunosuppression is particularly dependent on Egr2 signaling. We will investigate whether Egr2 signaling contributes to pro-tumor effects of obesity reprogrammed macrophages in future studies.

OA is the most abundant monounsaturated fatty acid in the human diet and highly enriched in olive oil^47^. Olive oil is a main component of the Mediterranean diet, and consumption of olive oil is inversely related to cancer prevalence^48^. It is generally thought that the beneficial effects of olive oil are due to its high proportion of OA, and studies have suggested that OA has potential anti-tumor effects^35,49^. However, prior studies reported that OA treatment induced M2 macrophage polarization^38,50^, which seems conflict with clinical observations and anti-tumor effects^37,51^. We have made a unique observation that SA and OA differentially regulate Egr2 expression in macrophages. OA can significantly abrogate SA-induced Egr2 expression and reverse SA-induced immunosuppression. These findings not only provide a mechanistic explanation for anti-tumor effects of OA but also provide a way to inhibit Egr2 expression and Egr2 mediated immunosuppression in macrophages. As Egr2 inhibitor is currently not available. The current work reveals a novel approach to inhibit Egr2-mediated immunosuppression by using OA. The data further supports the notion that diets enriched in OA may have potential for prevention and suppression of breast cancer.

## Acknowledgements

This work was partially supported by grants from NIH P20GM135004 (JY/CD), NIH R01CA278941 (JY), American Cancer Society MBGI-23-1150374-01-MBG (CD), NIH 1R21CA288604-01A1 (CD). IMC and CyTOF data acquisition and analysis were performed at UofL BCC Functional Immunomics Core supported by NIH P20GM135004.

## Reference List

1. Williams, C.B., Yeh, E.S., and Soloff, A.C. (2016). Tumor-associated macrophages: unwitting accomplices in breast cancer malignancy. npj Breast Cancer 2, 15025. 10.1038/npjbcancer.2015.25.

2. Garner, H., and de Visser, K.E. (2020). Immune crosstalk in cancer progression and metastatic spread: a complex conversation. Nature Reviews Immunology. 10.1038/s41577-019-0271-z.

3. Franklin, R.A., Liao, W., Sarkar, A., Kim, M.V., Bivona, M.R., Liu, K., Pamer, E.G., and Li, M.O. (2014). The cellular and molecular origin of tumor-associated macrophages. Science (New York, N.Y.) 344, 921–925. 10.1126/science.1252510.

4. Biswas, M. (2024). Understanding tissue-resident macrophages unlocks the potential for novel combinatorial strategies in breast cancer. Front Immunol 15, 1375528. 10.3389/fimmu.2024.1375528.

5. Harris, M.A., Savas, P., Virassamy, B., O’Malley, M.M.R., Kay, J., Mueller, S.N., Mackay, L.K., Salgado, R., and Loi, S. (2024). Towards targeting the breast cancer immune microenvironment. Nature Reviews Cancer 24, 554–577. 10.1038/s41568-024-00714-6.

6. Laviron, M., Petit, M., Weber-Delacroix, E., Combes, A.J., Arkal, A.R., Barthélémy, S., Courau, T., Hume, D.A., Combadière, C., Krummel, M.F., and Boissonnas, A. (2022). Tumor-associated macrophage heterogeneity is driven by tissue territories in breast cancer. Cell Reports 39, 110865. 10.1016/j.celrep.2022.110865.

7. Elfstrum, A.K., Bapat, A.S., and Schwertfeger, K.L. (2024). Defining and targeting macrophage heterogeneity in the mammary gland and breast cancer. Cancer Med 13, e7053. 10.1002/cam4.7053.

8. Hirano, R., Okamoto, K., Shinke, M., Sato, M., Watanabe, S., Watanabe, H., Kondoh, G., Kadonosono, T., and Kizaka-Kondoh, S. (2023). Tissue-resident macrophages are major tumor-associated macrophage resources, contributing to early TNBC development, recurrence, and metastases. Communications Biology 6, 144. 10.1038/s42003-023-04525-7.

9. Liu, Z., Gao, Z., Li, B., Li, J., Ou, Y., Yu, X., Zhang, Z., Liu, S., Fu, X., Jin, H., et al. (2022). Lipid-associated macrophages in the tumor-adipose microenvironment facilitate b east cancer progression. OncoImmunology 11, 2085432. 10.1080/2162402X.2022.2085432.

10. Timperi, E., Gueguen, P., Molgora, M., Magagna, I., Kieffer, Y., Lopez-Lastra, S., Sirven, P., Baudrin, L.G., Baulande, S., Nicolas, A., et al. (2022). Lipid-Associated Macrophages Are Induced by Cancer-Associated Fibroblasts and Mediate Immune Suppression in Breast Cancer. Cancer Research 82, 3291–3306. 10.1158/0008-5472.Can-22-1427.

11. Vassiliou, E., and Farias-Pereira, R. (2023). Impact of Lipid Metabolism on Macrophage Polarization: Implications for Inflammation and Tumor Immunity. Int J Mol Sci 24. 10.3390/ijms241512032.

12. Sun, R., Lei, C., Xu, Z., Gu, X., Huang, L., Chen, L., Tan, Y., Peng, M., Yaddanapudi, K., Siskind, L., et al. (2024). Neutral ceramidase regulates breast cancer progression by metabolic programming of TREM2-associated macrophages. Nature Communications 15, 966. 10.1038/s41467-024-45084-7.

13. Zhang, S., Lv, K., Liu, Z., Zhao, R., and Li, F. (2024). Fatty acid metabolism of immune cells: a new target of tumour immunotherapy. Cell Death Discovery 10, 39. 10.1038/s41420-024-01807-9.

14. Koundouros, N., Nagiec, M.J., Bullen, N., Noch, E.K., Burgos-Barragan, G., Li, Z., He, L., Cho, S., Parang, B., Leone, D., et al. (2025). Direct sensing of dietary ω-6 linoleic acid through FABP5-mTORC1 signaling. Science 387, eadm9805. doi:10.1126/science.adm9805.

15. Shao, N., Qiu, H., Liu, J., Xiao, D., Zhao, J., Chen, C., Wan, J., Guo, M., Liang, G., Zhao, X., and Xu, L. (2025). Targeting lipid metabolism of macrophages: A new strategy for tumor therapy. Journal of Advanced Research 68, 99–114. 10.1016/j.jare.2024.02.009.

16. Wan, M., Pan, S., Shan, B., Diao, H., Jin, H., Wang, Z., Wang, W., Han, S., Liu, W., He, J., et al. (2025). Lipid metabolic reprograming: the unsung hero in breast cancer progression and tumor microenvironment. Molecular Cancer 24, 61. 10.1186/s12943-025-02258-1.

17. Yu, L., Liebenberg, K., Shen, Y., Liu, F., Xu, Z., Hao, X., Wu, L., Zhang, W., Chan, H.L., Wei, B., et al. (2025). Tumor-derived arachidonic acid reprograms neutrophils to promote immune suppression and therapy resistance in triple-negative breast cancer. Immunity 58, 909–925.e907. 10.1016/j.immuni.2025.03.002.

18. Avellino, A., Jiang, X., Lee, M., Yu, J., Liu, S., Han, X., Li, J., Shilyansky, J., Wang, Z., Curry, M., et al. (2025). An Olive Oil–Based High-Fat Diet Promotes Obesity-Driven Metastasis of Triple-Negative Breast Cancer. Cancer Research 85, 5015–5032. 10.1158/0008-5472.Can-25-0822.

19. Williams, J.B., Horton, B.L., Zheng, Y., Duan, Y., Powell, J.D., and Gajewski, T.F. (2017). The EGR2 targets LAG-3 and 4-1BB describe and regulate dysfunctional antigen-specific CD8+ T cells in the tumor microenvironment. The Journal of experimental medicine 214, 381–400. 10.1084/jem.20160485.

20. Symonds, A.L.J., Miao, T., Busharat, Z., Li, S., and Wang, P. (2023). Egr2 and 3 maintain anti-tumour responses of exhausted tumour infiltrating CD8 + T cells. Cancer Immunol Immunother 72, 1139–1151. 10.1007/s00262-022-03319-w.

21. Wagle, M.V., Vervoort, S.J., Kelly, M.J., Van Der Byl, W., Peters, T.J., Martin, B.P., Martelotto, L.G., Nüssing, S., Ramsbottom, K.M., Torpy, J.R., et al. (2021). Antigen-driven EGR2 expression is required for exhausted CD8+ T cell stability and maintenance. Nature Communications 12, 2782. 10.1038/s41467-021-23044-9.

22. Zhang, Y., Li, H., Hao, Y., Chen, J., Chen, X., and Yin, H. (2025). EGR2 O-GlcNAcylation orchestrates the development of protumoral macrophages to limit CD8+ T cell antitumor responses. Cell Chemical Biology 32, 809–825.e807. 10.1016/j.chembiol.2025.05.007.

23. Wei, S., Xu, G., Zhao, S., Zhang, C., Feng, Y., Yang, W., Lu, R., Zhou, J., and Ma, Y. (2025). EGR2 promotes liver cancer metastasis by enhancing IL-8 expression through transcription regulation of PDK4 in M2 macrophages. International Immunopharmacology 153, 114484. 10.1016/j.intimp.2025.114484.

24. Ding, C., Sun, X., Wu, C., Hu, X., Zhang, H.-g., and Yan, J. (2019). Tumor Microenvironment Modulates Immunological Outcomes of Myeloid Cells with mTORC1 Disruption. The Journal of Immunology 202, 1623–1634. 10.4049/jimmunol.1801112.

25. Wu, C., Zhong, Q., Shrestha, R., Wang, J., Hu, X., Li, H., Rouchka, E.C., Yan, J., and Ding, C. (2023). Reactive myelopoiesis and FX-expressing macrophages triggered by chemotherapy promote cancer lung metastasis. JCI Insight 8. 10.1172/jci.insight.167499.

26. Wu, S.Z., Al-Eryani, G., Roden, D.L., Junankar, S., Harvey, K., Andersson, A., Thennavan, A., Wang, C., Torpy, J.R., Bartonicek, N., et al. (2021). A single-cell and spatially resolved atlas of human breast cancers. Nature Genetics 53, 1334–1347. 10.1038/s41588-021-00911-1.

27. Cui, H., Zhao, G., Lu, Y., Zuo, S., Duan, D., Luo, X., Zhao, H., Li, J., Zeng, Z., Chen, Q., and Li, T. (2025). TIMER3: an enhanced resource for tumor immune analysis. Nucleic Acids Research 53, W534–W541. 10.1093/nar/gkaf388.

28. Jablonski, K.A., Amici, S.A., Webb, L.M., Ruiz-Rosado, J.d.D., Popovich, P.G., Partida-Sanchez, S., and Guerau-de-Arellano, M. (2015). Novel Markers to Delineate Murine M1 and M2 Macrophages. PloS one 10, e0145342–e0145342. 10.1371/journal.pone.0145342.

29. Veremeyko, T., Yung, A.W.Y., Anthony, D.C., Strekalova, T., and Ponomarev, E.D. (2018). Early Growth Response Gene-2 Is Essential for M1 and M2 Macrophage Activation and Plasticity by Modulation of the Transcription Factor CEBPβ. Frontiers in immunology 9, 2515–2515. 10.3389/fimmu.2018.02515.

30. Tu, H.-F., Tao, J., Hu, M.-H., Fan, D., Tsai, Y.-C., Wu, T.-C., and Hung, C.-F. (2025). Type I Interferon Modulates the Function of Ly6C High-Expressing Naïve CD8+ T Cells to Promote an Antitumor Response. Vaccines 13, 246.

31. Riquelme, P., Amodio, G., Macedo, C., Moreau, A., Obermajer, N., Brochhausen, C., Ahrens, N., Kekarainen, T., Fändrich, F., Cuturi, C., et al. (2017). DHRS9 Is a Stable Marker of Human Regulatory Macrophages. Transplantation 101.

32. Devalaraja, S., To, T.K.J., Folkert, I.W., Natesan, R., Alam, M.Z., Li, M., Tada, Y., Budagyan, K., Dang, M.T., Zhai, L., et al. (2020). Tumor-Derived Retinoic Acid Regulates Intratumoral Monocyte Differentiation to Promote Immune Suppression. Cell 180, 1098–1114.e1016. 10.1016/j.cell.2020.02.042.

33. Figley, M.D., Gu, W., Nanson, J.D., Shi, Y., Sasaki, Y., Cunnea, K., Malde, A.K., Jia, X., Luo, Z., Saikot, F.K., et al. (2021). SARM1 is a metabolic sensor activated by an increased NMN/NAD+ ratio to trigger axon degeneration. Neuron 109, 1118–1136.e1111. 10.1016/j.neuron.2021.02.009.

34. Cameron, A.M., Castoldi, A., Sanin, D.E., Flachsmann, L.J., Field, C.S., Puleston, D.J., Kyle, R.L., Patterson, A.E., Hässler, F., Buescher, J.M., et al. (2019). Inflammatory macrophage dependence on NAD+ salvage is a consequence of reactive oxygen species–mediated DNA damage. Nature Immunology 20, 420–432. 10.1038/s41590-019-0336-y.

35. Kim, J.S., Kim, D.K., Moon, J.Y., Lee, M.-Y., and Cho, S.K. (2024). Oleic acid inhibits the migration and invasion of breast cancer cells with stemness characteristics through oxidative stress-mediated attenuation of the FAK/AKT/NF-κB pathway. Journal of Functional Foods 116, 106224. 10.1016/j.jff.2024.106224.

36. Deng, B., Kong, W., Suo, H., Shen, X., Newton, M.A., Burkett, W.C., Zhao, Z., John, C., Sun, W., Zhang, X., et al. (2023). Oleic Acid Exhibits Anti-Proliferative and Anti-Invasive Activities via the PTEN/AKT/mTOR Pathway in Endometrial Cancer. Cancers (Basel) 15. 10.3390/cancers15225407.

37. Zhang, Y., Xiang, Z., Xu, Y., Cheung, L.S., Wang, X., Wang, M., Wong, H.H.W., Zhu, Z., Zhang, W., Gao, Y., et al. (2025). Oleic acid restores the impaired antitumor immunity of γδ-T cells induced by palmitic acid. Signal Transduction and Targeted Therapy 10, 209. 10.1038/s41392-025-02295-8.

38. Hou, Y., Wei, D., Zhang, Z., Guo, H., Li, S., Zhang, J., Zhang, P., Zhang, L., and Zhao, Y. (2022). FABP5 controls macrophage alternative activation and allergic asthma by selectively programming long-chain unsaturated fatty acid metabolism. Cell Reports 41. 10.1016/j.celrep.2022.111668.

39. Iwata, A., Maruyama, J., Natsuki, S., Nishiyama, A., Tamura, T., Tanaka, M., Shichino, S., Seki, T., Komai, T., Okamura, T., et al. (2024). Egr2 drives the differentiation of Ly6Chi monocytes into fibrosis-promoting macrophages in metabolic dysfunction-associated steatohepatitis in mice. Communications Biology 7, 681. 10.1038/s42003-024-06357-5.

40. Shao, N., Qiu, H., Liu, J., Xiao, D., Zhao, J., Chen, C., Wan, J., Guo, M., Liang, G., Zhao, X., and Xu, L. (2024). Targeting lipid metabolism of macrophages: A new strategy for tumor therapy. Journal of Advanced Research. 10.1016/j.jare.2024.02.009.

41. Kunkemoeller, B., Prendeville, H., McIntyre, C., Temesgen, A., Loftus, R.M., Yao, C., Dyck, L., Sinclair, L.V., Rollings, C., Douglas, A., et al. (2025). The source of dietary fat influences anti-tumour immunity in obese mice. Nature Metabolism 7, 1630–1645. 10.1038/s42255-025-01330-w.

42. LeBlanc, S.E., Srinivasan, R., Ferri, C., Mager, G.M., Gillian-Daniel, A.L., Wrabetz, L., and Svaren, J. (2005). Regulation of cholesterol/lipid biosynthetic genes by Egr2/Krox20 during peripheral nerve myelination. Journal of Neurochemistry 93, 737–748. 10.1111/j.1471-4159.2005.03056.x.

43. Iwata, A., Maruyama, J., Natsuki, S., Nishiyama, A., Tamura, T., Tanaka, M., Shichino, S., Seki, T., Komai, T., Okamura, T., et al. (2024). Egr2 drives the differentiation of Ly6C(hi) monocytes into fibrosis-promoting macrophages in metabolic dysfunction-associated steatohepatitis in mice. Commun Biol 7, 681. 10.1038/s42003-024-06357-5.

44. Angela, M., Endo, Y., Asou, H.K., Yamamoto, T., Tumes, D.J., Tokuyama, H., Yokote, K., and Nakayama, T. (2016). Fatty acid metabolic reprogramming via mTOR-mediated inductions of PPARγ directs early activation of T cells. Nature Communications 7, 13683. 10.1038/ncomms13683.

45. Chowdhury, P.S., Chamoto, K., Kumar, A., and Honjo, T. (2018). PPAR-Induced Fatty Acid Oxidation in T Cells Increases the Number of Tumor-Reactive CD8+ T Cells and Facilitates Anti–PD-1 Therapy. Cancer Immunology Research 6, 1375–1387. 10.1158/2326-6066.Cir-18-0095.

46. Tiwari, P., Blank, A., Cui, C., Schoenfelt, K.Q., Zhou, G., Xu, Y., Khramtsova, G., Olopade, F., Shah, A.M., Khan, S.A., et al. (2019). Metabolically activated adipose tissue macrophages link obesity to triple-negative breast cancer. The Journal of experimental medicine 216, 1345–1358. 10.1084/jem.20181616.

47. Di Serio, M.G., Di Giacinto, L., Di Loreto, G., Giansante, L., Pellegrino, M., Vito, R., and Perri, E. (2016). Chemical and sensory characteristics of Italian virgin olive oils from Grossa di Gerace cv. European Journal of Lipid Science and Technology 118, 288–298. 10.1002/ejlt.201400622.

48. Psaltopoulou, T., Kosti, R.I., Haidopoulos, D., Dimopoulos, M., and Panagiotakos, D.B. (2011). Olive oil intake is inversely related to cancer prevalence: a systematic review and a meta-analysis of 13800 patients and 23340 controls in 19 observational studies. Lipids in Health and Disease 10, 127. 10.1186/1476-511X-10-127.

49. Pascual, G., Domínguez, D., Elosúa-Bayes, M., Beckedorff, F., Laudanna, C., Bigas, C., Douillet, D., Greco, C., Symeonidi, A., Hernández, I., et al. (2021). Dietary palmitic acid promotes a prometastatic memory via Schwann cells. Nature 599, 485–490. 10.1038/s41586-021-04075-0.

50. Cao, J., Chen, K., Hu, K., Mi, X., Pan, Y., Xiao, D., Liu, S., Xiao, L., Zhou, L., Tao, Y., and Tang, J. (2025). USP2-mediated PPARγ stabilization promotes hepatocellular carcinoma progression and M2 macrophage polarization via oleic acid. Journal for ImmunoTherapy of Cancer 13, e012721. 10.1136/jitc-2025-012721.

51. Prendeville, H., and Lynch, L. (2022). Diet, lipids, and antitumor immunity. Cellular & Molecular Immunology 19, 432–444. 10.1038/s41423-021-00781-x.

