## Supplemental data for "Tumor-derived stearic acid induces macrophage Egr2 signaling to suppress anti-tumor immunity in breast cancer"

**Supplemental Table S1.** Antibodies used in Flow cytometry and Western blotting

| **Antibodies** | **Company and catalog number** |
|---|---|
| PerCP/Cyanine5.5 anti-mouse CD45 | BioLegend, No. 157208 |
| PE/Cyanine7 anti-mouse CD45 | BioLegend, No. 103114 |
| APC anti-mouse/human CD11b | BioLegend, No. 101212 |
| PE anti-mouse Ly6G | BioLegend, No. 127608 |
| PerCP anti-mouse Ly6C | BioLegend, No. 128028 |
| PE anti-mouse F4/80 | BioLegend, No. 123110 |
| APC anti-mouse CD4 | BioLegend, No. 100412 |
| FITC anti-mouse CD8a | BioLegend, No. 100706 |
| APC anti-mouse NK-1.1 | BioLegend, No. 108710 |
| PE-anti-mouse TNFα | BioLegend, No. 506306 |
| PE anti-mouse IFN-γ | BioLegend, No. 505808 |
| PE-Cy7 anti-mouse IFN-γ | BioLegend, No. 505826 |
| FITC anti-mouse I-A/I-E | BioLegend, No. 107606 |
| APC anti-mouse Egr2 | Invitrogen, No. 17-6691-82 |
| Fixable Viability Dye eFluor™ 780 | Thermo Fisher Scientific, No. 65-0865-14 |
| Anti-mouse PPARγ | Cell Signaling, No. 2443 |

**Supplemental Table S2.** Antibodies used in CyTOF and IMC

| | **Antibodies** | **Company and catalog number** |
|---|---|---|
| 1 | Anti-Mouse CD45 (30-F11)-89Y | Fluidigm, No. 3089005B |
| 2 | Anti-Mouse CD11c (N418)-142Nd | Fluidigm, No. 3142003B |
| 3 | Anti-Mouse CD69 (H1.2F3)-143Nd | Fluidigm, No. 3143004B |
| 4 | Anti-Mouse CD4 (RM4-5)-145Nd | Fluidigm, No. 3145002B |
| 5 | Anti-Mouse F4/80 (BM8)-146Nd | Fluidigm, No. 3146008B |
| 6 | Anti-Mouse CD103 (2E7)-148Nd | Biolegend, No. 121402 |
| 7 | Anti-Mouse CD19 (6D5)-149Sm | Fluidigm, No. 3149002B |
| 8 | Anti-Mouse Ly-6C (HK1.4)-150Nd | Fluidigm, No. 3150010B |
| 9 | Anti-Mouse CD25 (3C7)-151Eu | Fluidigm, No. 3151007B |
| 10 | Anti-Mouse CD3e (145-2C11)-152Sm | Fluidigm, No. 3152004B |
| 11 | Anti-Mouse CD274/PD-L1-153Eu (10F.9G2) | Fluidigm, No. 3153016B |
| 12 | Anti-Mouse PD-1 (29F.1A12)-159Tb | Fluidigm, No. 3159024B |
| 13 | Anti-Mouse CD62L (MEL-14)-160Gd | Fluidigm, No. 3160008B |
| 14 | Anti-Human/Mouse CD44 (IM7)-162Dy | Fluidigm, No. 3162030B |
| 15 | Anti-Mouse CX3CR1 (SA011F11)-164Dy | Fluidigm, No. 3164023B |
| 16 | Anti-Mouse CD8a (53-6.7)-168Er | Fluidigm, No. 3168003B |
| 17 | Anti-Mouse CD206/MMR (C068C2)-169Tm | Fluidigm, No. 3169021B |
| 18 | Anti-Mouse NK1.1 (PK136)-170Er | Fluidigm, No. 3170002B |
| 19 | Anti-Mouse CD11b (M1/70 )-172Yb | Fluidigm, No. 3172012B |
| 20 | Anti-Mouse CD223/LAG3 (C9B7W)-174Yb | Fluidigm, No. 3174019B |
| 21 | Anti-Human/Mouse CD45R/B220 (RA3-6B2)-176Yb | Fluidigm, No. 3176002B |
| 22 | Anti-Mouse I-A/I-E (M5/114.15.2)-209Bi | Fluidigm, No. 3209006B |
| 23 | Anti-Mouse CD127/IL7Ra (A7R34)-175Lu | Fluidigm, No. 3175006B |
| 24 | Anti-Mouse iNOS (CXNFT)-161Dy | Fluidigm, No. 3161011B |
| 24 | Anti-Mouse TNFα (MP6-XT22)-141Pr | Fluidigm, No. 3141013B |
| 25 | Anti-Mouse IL-2 (JES6-5H4)-144Nd | Fluidigm, No. 3144002B |
| 26 | Anti-Mouse CCR2 (475301R)-156Gd | R&D System, No. MAB55381R |
| 27 | Anti-Mouse IFNg (XMG1.2)-165Ho | Fluidigm, No. 3165003B |
| 28 | Anti-Mouse IL-6 (MP5-20F3)-167Er | Fluidigm, No. 3167003B |
| 29 | Anti-Mouse Foxp3 (FLK-16s)-158Gd | Fluidigm, No. 3158003A |
| 30 | Anti-Human CD8a (C8/144B)-162Dy | Standard BioTools, No. 3162034D |
| 31 | Anti-Human CD68 (KP1)-141Pr | Standard BioTools, No. 91H012141 |
| 32 | Anti-Egr2 | GeneTex, No. GTX102912 |

**Supplemental Table S3.** Primer sequences for real-time PCR

| Gene | Forward primer | Reverse primer |
|---|---|---|
| CEBPB | ACTTCAGCCCCTACCTGGAG | GGCTCACGTAACCGTAGTCG |
| PPARG | TGTCGGTTTCAGAAGTGCCT | CCAACAGCTTCTCCTTCTCG |
| CHIL3 | ACTTTGATGGCCTCAACCTG | AATGATTCCTGCTCCTGTGG |
| ARG1 | TTTTAGGGTTACGGCCGGTG | CCTCGAGGCTGTCCTTTTGA |
| RETNLA | CTCATCTGCATCTCCCTGCT | AGGAGGCCCATCTGTTCATAG |

**Supplemental Figure S1**

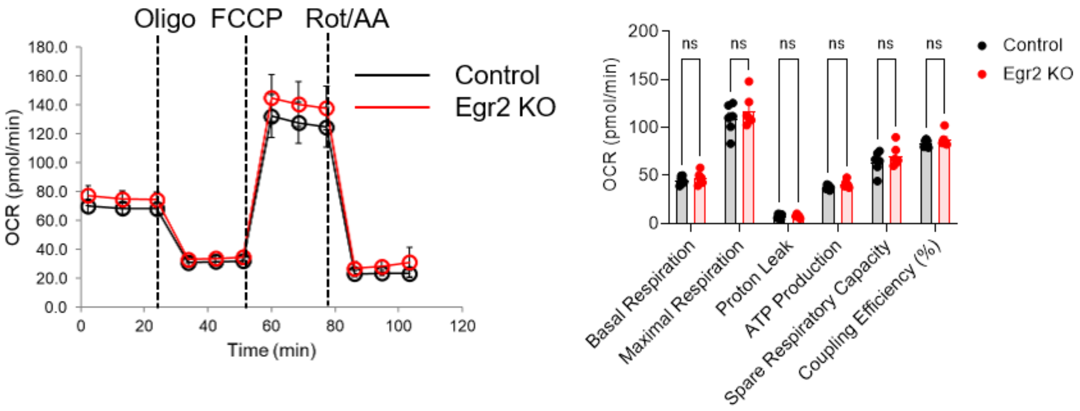


**Figure S1.** Seahorse Mito stress test in TAMs from E0771 tumors of control and Egr2 KO mice. Summarized of relative values of ECAR bioenergenic profiling was shown. ns: not significant.

**Supplemental Figure S2**

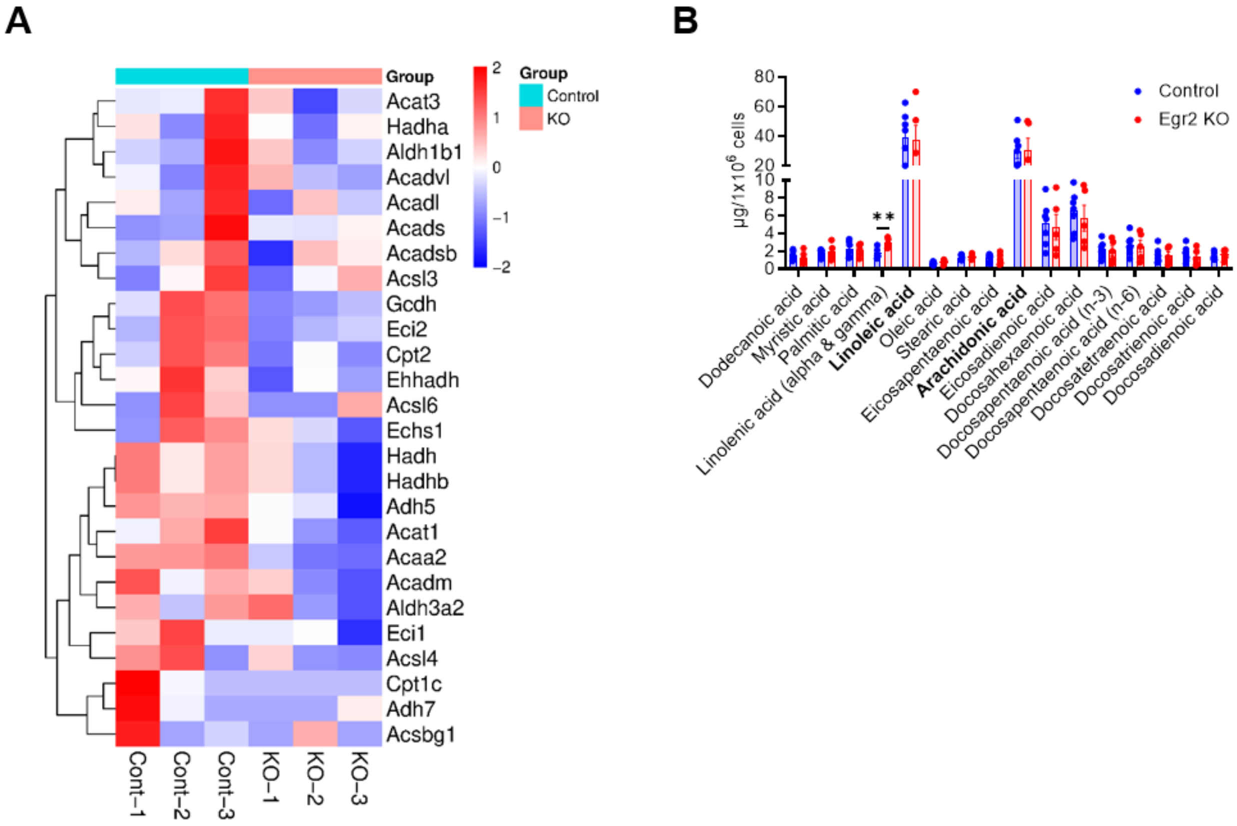


**Figure S2.** (A) Heatmap showing the clustering of fatty acid degradation related genes in TAMs from E0771 tumor-bearing control and Egr2 KO mice (n=3) based on log-relative abundances. (B) Contents and levels of fatty acid in the lysates of TAMs from E0771 tumor-bearing control and Egr2 KO mice.

**Supplemental Figure S3**

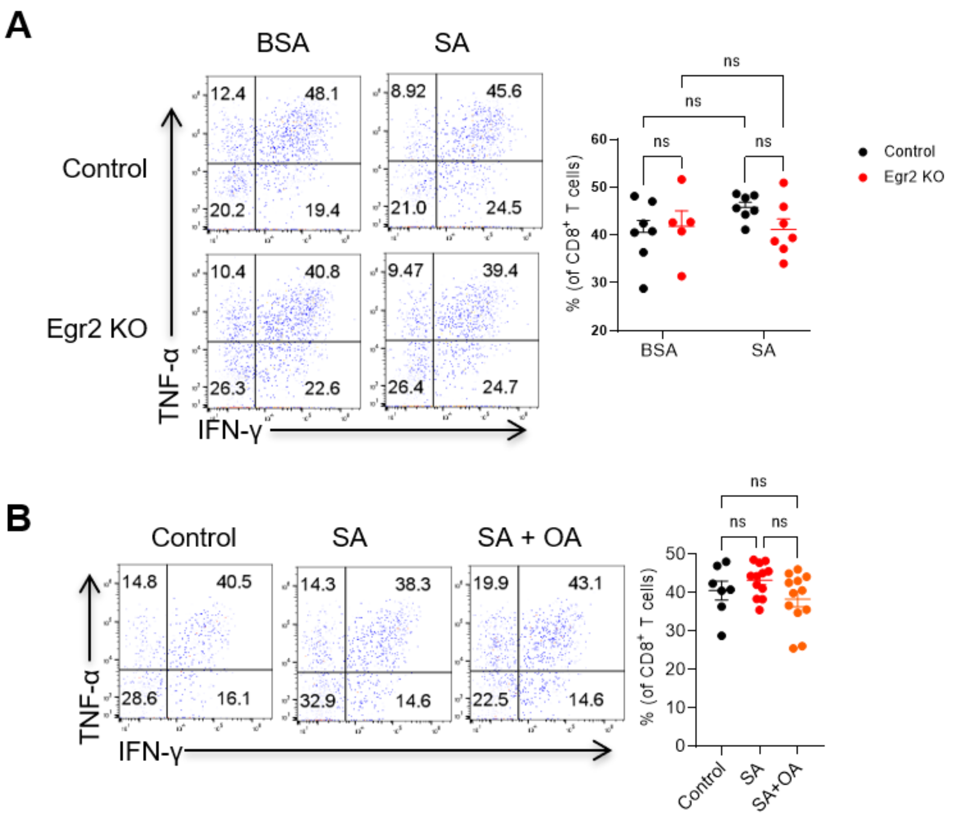


**Figure S3.** (A) Percentages of IFN-γ$^+$TNF-α$^+$ CD8$^+$ T cells on day 7 in tumor cell and macrophage admix experiments in 4 groups**.** (B) Percentages of IFN-γ$^+$TNF-α$^+$ CD8$^+$ T cells on day 7 in tumor cell and macrophage admix experiments in 3 groups. Each dot represents an individual mouse. ns: not significant.
